# Optogenetic stimulation of BLA terminals in the BNST elicits long term synaptic plasticity and restores synaptic alterations induced by adolescent *Shank3* downregulation

**DOI:** 10.1101/2021.04.02.438185

**Authors:** Christelle Glangetas, Alessandro Contestabile, Pedro Espinosa, Giulia Casarotto, Stefano Musardo, Camilla Bellone

**Author notes:** These authors contributed equally to this work (Christelle Glangetas, Alessandro Contestabile).

## Abstract

Anxiety disorders are the most prevalent co-morbidity factor associated with the core domains of Autism Spectrum Disorders (ASD). Investigations on potential common neuronal mechanisms that may explain the co-occurrence of ASD and anxiety disorders are still poorly unexplored. One of the key questions which remained unsolved is the role of Shank3 protein in anxiety behaviors. Here we used shRNA strategy to model Shank3 insufficiency in the bed nucleus of the stria terminalis (BNST). We found that *Shank3* downregulation in the BNST induced anxiogenic effects. Associated with these behavioural defects, we showed alteration of glutamatergic synaptic functions in the BNST induced by S*hank*3 insufficiency during adolescence. In addition, we pointed that adolescence represents a crucial time window to interfere with BNST maturation, a key hub for anxiety control. Together, these results unravelled the crucial role of S*hank*3 expression in the BNST during adolescence. Our study provided a new insight in the neuronal mechanisms underlying anxiety disorders. We proposed to screen a novel molecular target essential for BNST integrity during adolescence. This result may further help for the diagnosis and the development of therapeutic strategy for anxiety disorders and anxiety disorders implicated in some forms of ASD.

## Introduction

Autism Spectrum disorders (ASD) which estimated prevalence is around 1 in 68 people have been characterized by a triad of symptoms: deficit in social interactions and communications with repetitive or obsessive compulsive behaviors. Anxiety disorders are among the most frequent comorbidity factors associated with these core domains which can affect up to 40 percent of ASD patients^1,2^. Anxiety is a physiological response which helps to face a potential threat and to promote survival but if persistent can be maladaptive and further turns into anxiety disorders. According to the DSM-V, anxiety disorders can be classified into distinct categories based on the focus of anxiety such as separation anxiety disorder, panic disorder, specific phobia, social anxiety disorder, agoraphobia or generalized anxiety disorder. One remaining question which is still unsolved is to determine why ASD children have a higher prevalence in their life to develop anxiety disorders compared to typical developing children^3–5^. It has been shown that anxiety disorders can alter the quality of life in typical children and may exacerbate ASD symptoms ^6,7^. Investigations on the links and potential neurobiological mechanisms that may explain the co-occurrence of ASD and anxiety disorders represent a crucial challenge. These studies may help to refine the development of specific therapeutic strategies for anxiety disorders in ASD children which are still needed.

Despite the complexity of ASD ethology, there is a strong genetic component in these disorders. In particular, mutation of excitatory post synaptic scaffolding proteins like SHANK3 has been identified as responsible of some ASD forms like in the Phelan-McDermid Syndrome in which anxiety disorder has also been reported^8–13^. In addition, some mouse models generated to mimic ASD also present anxiety traits as in *Shank3* mutant mice^14–17^. Interestingly, the peak in anxiety phenotype is notably reported between school age children and adolescence in ASD patients^18–21^. Adolescence which is approximately between P28 and P55 in mice could appear as a crucial time window for the maturation of anxiety circuits ^22,23^ and interfering with these circuits during this critical developmental window may have behavioral consequences. Despite re-expression of *Shank3* gene in adult *Shank3* knock-in mice restored social deficits and grooming alterations, anxiety traits still remained^24^. Interestingly, anxiety was rescued when *Shank3* was re-expressed during adolescence suggesting that *Shank3* gene expression in adolescence might be essential for anxiety circuit development. Understanding which neuronal circuits drive anxiety and how they mature are still open questions.

The Bed Nucleus of the Stria Terminalis (BNST) has been recognized as a key structure in exerting a powerful control on anxiety level both in humans and rodents^25–30^. Accumulative evidences point out a link between altered BNST activation and anxiety disorders in humans^31–36^. Moreover, BNST receives strong excitatory glutamatergic innervations notably from ventral subiculum and the basolateral amygdala which are both important in anxiety control^29,30,37,38^. Thus, we hypothesized that *Shank3* deficiency in the BNST during adolescence may affect the maturation of its excitatory synapses and therefore may trigger an anxiety phenotype in mice.

Here, to evaluate the consequences of *Shank3* gene downregulation in the BNST, we used a combination of cutting-edge technologies and anxiety behavioural assays associated with *Shank3* gene silencing strategy. We demonstrate that i) selective deficiency of *Shank3* in the BNST triggers anxiety-related behaviour in adolescent mice, ii) BNST *Shank3* insufficiency affects its excitatory synapses and iii) *in vivo* long-term depression of BLA to BNST synapses restores these synaptic alterations. Altogether, our study provides a new insight in the effect of *Shank3* deficiency on anxiety circuit maturation during adolescence in mice. This project highlights a novel molecular and circuit targets which may help for the development of future therapies for anxiety disorders.

## Results

### *Shank3* deficient mice present social behavior impairments and an anxiety phenotype in early adolescence

It is well known that human *SHANK3* gene codes for a scaffolding protein located in excitatory synapses and its haplo-insufficency is acknowledged to lead to a severe form of ASD, known as Phelan-McDermid syndrome^39,40^. Patients affected with this syndrome mainly present difficulties with social communication and interactions, deficitary locomotion, anxiety disorders and repetitive and/or obsessive compulsive behaviors^9^. In order to translationally study the Phelan-McDermid syndrome, rodent models with *Shank3* deletion have been implemented. Remarkably, adult male mice with a homozygote *Shank3*^Δe4-22^ mutation (*Shank3*^−/−^) presented a decreased interaction time with a juvenile sex-matched conspecific compared to wildtype littermate (*Shank3*^+/+^, **Sup. Fig. 1A-B**). Moreover, *Shank3*^−/−^ adult mice showed a decreased distance moved during the open field test (**Sup. Fig. 1C-D**) and an anxiety phenotype in both the open field test (**Sup. Fig. 1E**) and in the elevated plus maze (**Sup. Fig. 1F-I**). These results suggest that mice with a *Shank3*^Δe4-22^ mutation present neurological deficits similar to patients affected by Phelan-McDermid syndrome proving the relevance of this model in translational research.

We initially used *Shank3*^Δe4-22^ mutated mice to perform longitudinal analysis of the behavioral phenotypes and examine the development and the appearance of the behavioral deficits. For this purpose, we followed male and female mice generated by *Shank3*^+/−^ parents (**Fig. 1A-B**) in a longitudinal study where each 10 days we measured social interaction, locomotion and anxious behavior starting from P16 and ending at P56 (**Fig. 1C-E**). In the free interaction test, *Shank3*^−/−^ showed a decreased time interaction compared to *Shank3*^+/+^ control mice starting from P26 (**Fig. 1F-G**). Consistently with previous data, social deficits of *Shank3*^−/−^ mice were observed until adulthood (**Fig. 1F-G**). In parallel, distance travelled in an open field test was not significantly different between *Shank3*^−/−^ and *Shank3*^+/+^ control mice suggesting that deficit in locomotion may emerge later in life (**Fig. 1H-I**). Remarkably, *Shank3* deficient mice decreased the time spent in the center of the open field arena starting from P36 compared to control littermates suggesting that anxiety phenotype already emerged during adolescence (**Fig. 1J-K**).

**Figure 1 -.**
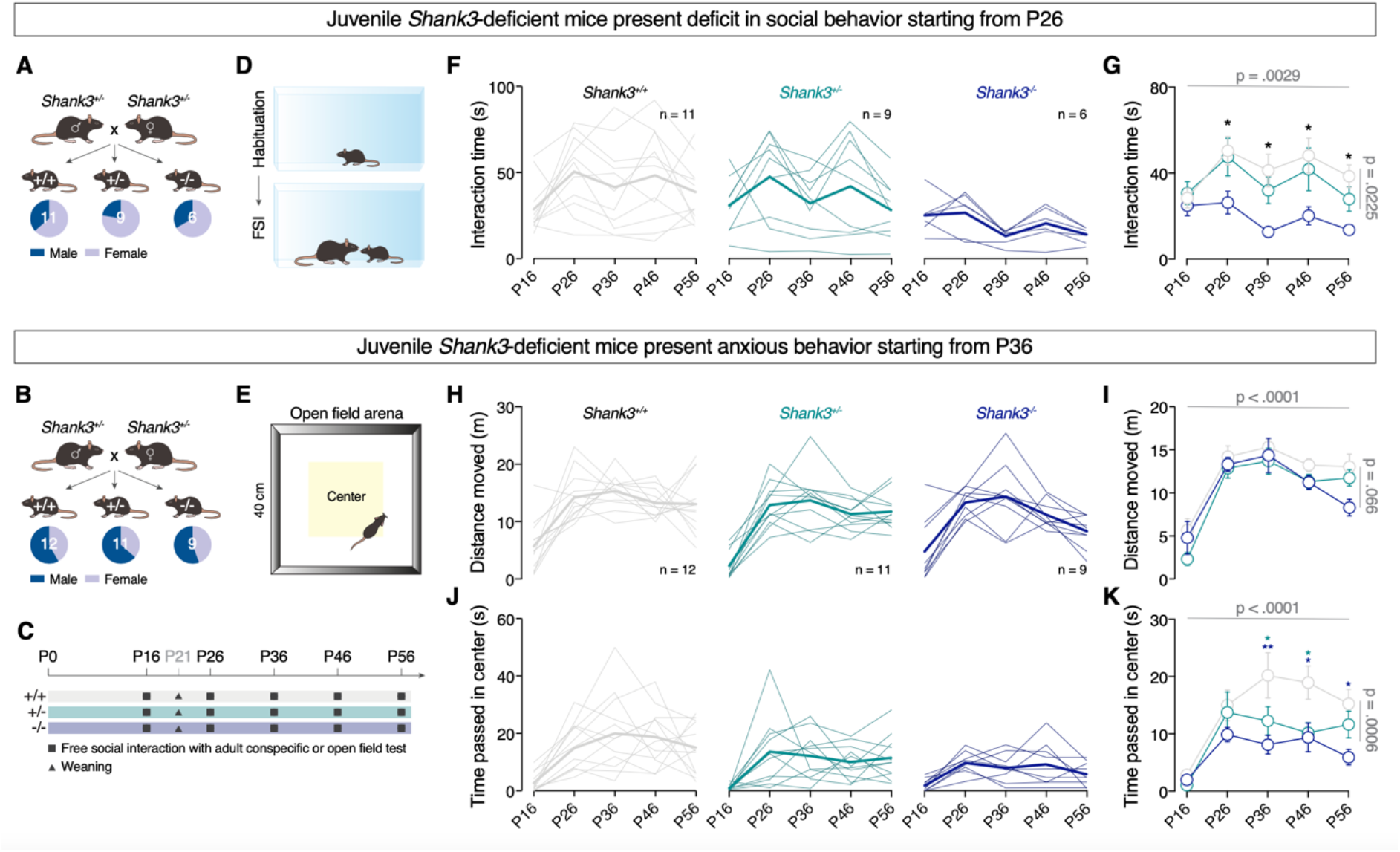
Development of social and exploring deficits and anxiety-like behaviour in *Shank3*^Δe4-22^ mutated mice. (**A** and **B**) proportion of male and female juvenile mice that performed the free social interaction (A) and open field test (B). (**C**) Longitudinal timeline. (**D** and **E**) Schematic representation of the free social interaction (D) and open field test (E). (**F** and **G**) Kinetic of the interaction time (s) in the free social interaction test across ages displayed by *Shank3*^+/+^, *Shank3*^+/−^ or *Shank3*^−/−^ mice expressed per intragroup (F) or intergroups (G: two-way RM ANOVA followed by Bonferroni’s multiple comparisons post-hoc test. Genotype x Time interaction: F_(8, 92)_ = 1.008, P = 0.4359; Time main effect: F_(4, 92)_ = 4.351, P = 0.0029; Genotype main effect: F_(2, 23)_ = 4.498, P = 0.0225). (**H** and **I**) Kinetic of the distance moved (cm) in the open field test across ages displayed by *Shank3*^+/+^, *Shank3*^+/−^ or *Shank3*^−/−^ mice expressed per intragroup (H) or intergroups (I: two-way RM ANOVA followed by Bonferroni’s multiple comparisons post-hoc test. Genotype x Time interaction: F(8, 116) = 0.9938, P = 0.447; Time main effect: F_(4, 116)_ = 38.01, P < 0.0001; Genotype main effect: F_(2, 29)_ = 2.994, P = 0.0658). (**J** and **K**) Kinetic of the time passed in the center (s) in the open field test across ages displayed by *Shank3*^+/+^, *Shank3*^+/−^ or *Shank3*^−/−^ mice expressed per intragroup (J) or intergroups (K: two-way RM ANOVA followed by Bonferroni’s multiple comparisons post-hoc test. Genotype x Time interaction: F_(8, 116)_ = 1.187, P = 0.3128; Time main effect: F_(4, 116)_ = 12.62, P < 0.0001; Genotype main effect: F_(2, 29)_ = 9.582, P = 0.0006). Abbreviations: *p < 0.05; **0.05 < p < 0.01.

These data show that different behavioral phenotypes present different developmental trajectories in *Shank3* deficient mice. Social behavior impairments seem to appear immediately after weaning and the anxiety phenotype only 10 days after.

### *Shank3* deficiency in the BNST drives anxiety in adolescent mice

In a parallel study, we demonstrated that selective postnatal downregulation of *Shank3* in the Ventral tegmental area (VTA) alters the sociability of mice ^41^. In the same manner, we were interested in manipulating the *Shank3* expression of a brain region implicated in anxiety in order to mimic the phenotype observed in *Shank3*^−/−^ mice. Given the emerging importance of the Bed Nucleus of the Stria Terminalis (BNST) in anxiety processing ^25,36^, we focused our investigation on this brain region. Similarly to the study conducted by Bariselli and co-authors, we selectively downregulated the BNST expression of *Shank3* in the first days after weaning (P21-24) by bilaterally injecting a virus expressing a sh*Shank3* (AAV-shShank3-luczsGreen) (**Fig. 2A**). A scramble version of the virus was injected as control and the efficacy of the sh*Shank3* virus was verified through RT-qPCR at P36 (**Fig. 2B**). Localization of the viral infection was controlled post hoc (**Fig. 2C**).

**Figure 2 -.**
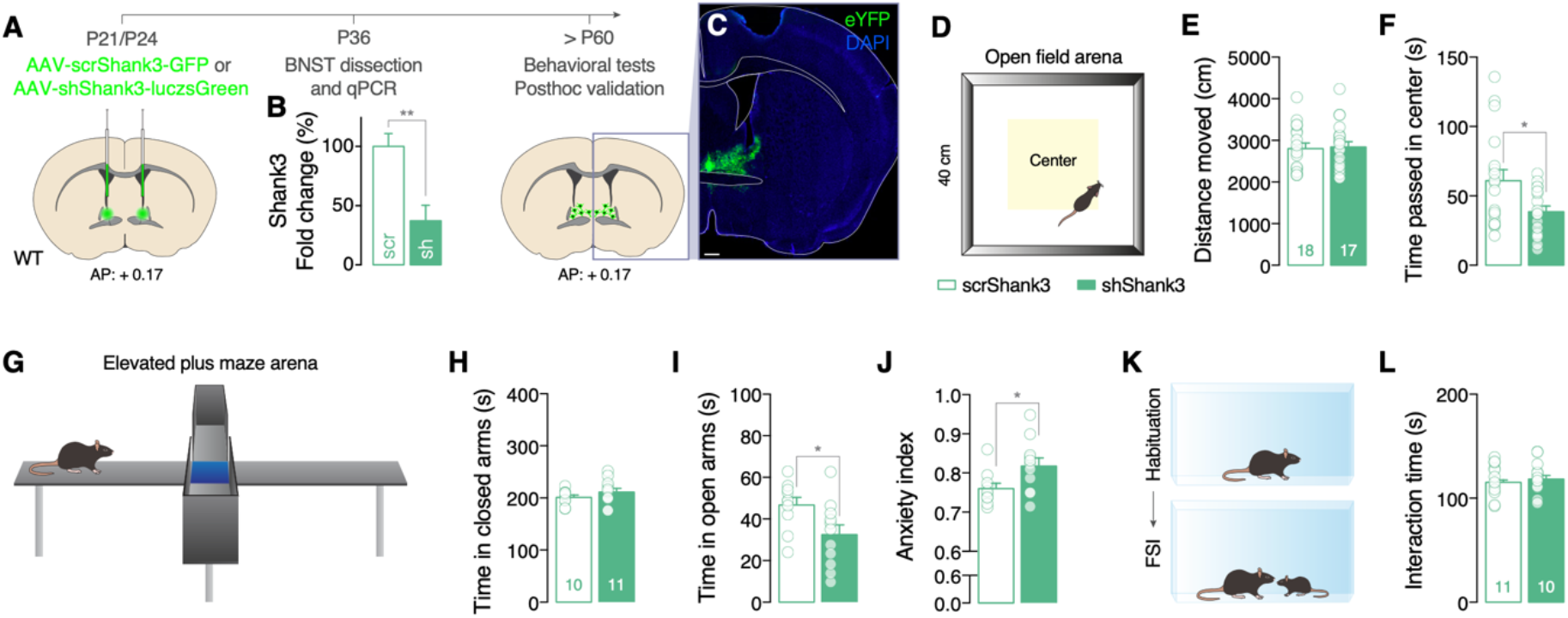
*Shank3* downregulation in the BNST is sufficient to trigger anxiety phenotype. (**A**) Experimental design. (**B**) RT-qPCR quantification of *Shank3* gene expression in the BNST of scr- or shShank3 injected mice. (**C**) Representative confocal image of a coronal slice of a brain injected with AAV-sh*Shank3*-luczsGreen in the BNST (scale bar = 500 μm). (**D, G** and **K**) Schematic representation of the open field (D), elevated plus maze (G) and free social interaction test (K). (**E** and **F**) Distance moved (E: unpaired t-test. t_(36)_ = 0.2006, P = 0.8421) and time passed in the center (F: Mann Whitney test. U = 90, P = 0.0380) during the open field test for scr- or shShank3 infected mice. (**H, I** and **J**) Time passed in the closed (H: unpaired t-test. t_(19)_ = 1.122, P = 0.2757) and open arms (I: unpaired t-test. t_(19)_ = 2.353, P = 0.0295) and calculated anxiety index (J: unpaired t-test. t_(19)_ = 2.315, P = 0.0320) in the elevated plus maze test for scr- or shShank3 infected mice. (**L**) Interaction time during free social interaction test for scr- or shShank3 infected mice (unpaired t-test. t_(19)_ = 0.1666, P = 0.916). Abbreviations: AP = antero-posterior axis; FSI = free social interaction; *p < 0.05.

In order to compare the behavior of *Shank3* deficient mice and the mice with downregulation of *Shank3* in the BNST, we performed the same battery of behavioral tests presented in **Sup. Fig. 1** during adulthood. Interestingly, no difference was observed between scr*Shank3-* and sh*Shank3-*expressing animals in the distance travelled during the open field (**Fig. 2D-E**). On the other hand, mice injected in the BNST with sh*Shank3* spent less time in the center of the open field suggesting an anxiety phenotype (**Fig. 2F**). Elevated plus maze test confirmed the anxiety phenotype for the sh*Shank3*-expressing animal since they passed less time in open arms (**Fig. 2G-I**) and their anxiety index was higher than the control group (**Fig. 2J**). Finally, the free social interaction test did not reveal any difference between the groups (**Fig. 2K-L**).

This data indicates that a downregulation of *Shank3* in the BNST during adolescence causes anxiety disorders without affecting social behavior and exploration of the open field arena.

### Optogenetic stimulation of BLA to BNST projections elicits long term synaptic plasticity and restores synaptic alterations induced by *Shank3* downregulation

After revealing the anxiety phenotype induced by *Shank3* downregulation in the BNST of adolescent mice, we evaluated the effect of this downregulation at the synaptic level. For this purpose, we injected a rAAV5-CaMKII-hChR2(H134R)-eYFP-WPRE virus in the basolateral amygdala (BLA) and we measured the light-evoked AMPA/NMDA ratio of scrShank3 and shShank3 positive neurons in the BNST (**Fig. 3A**). Interestingly, we found an increased AMPA/NMDA ratio in shShank3 expressing cells compared to control suggesting an alteration of the synapses composition (**Fig. 3B**).

**Figure 3 -.**
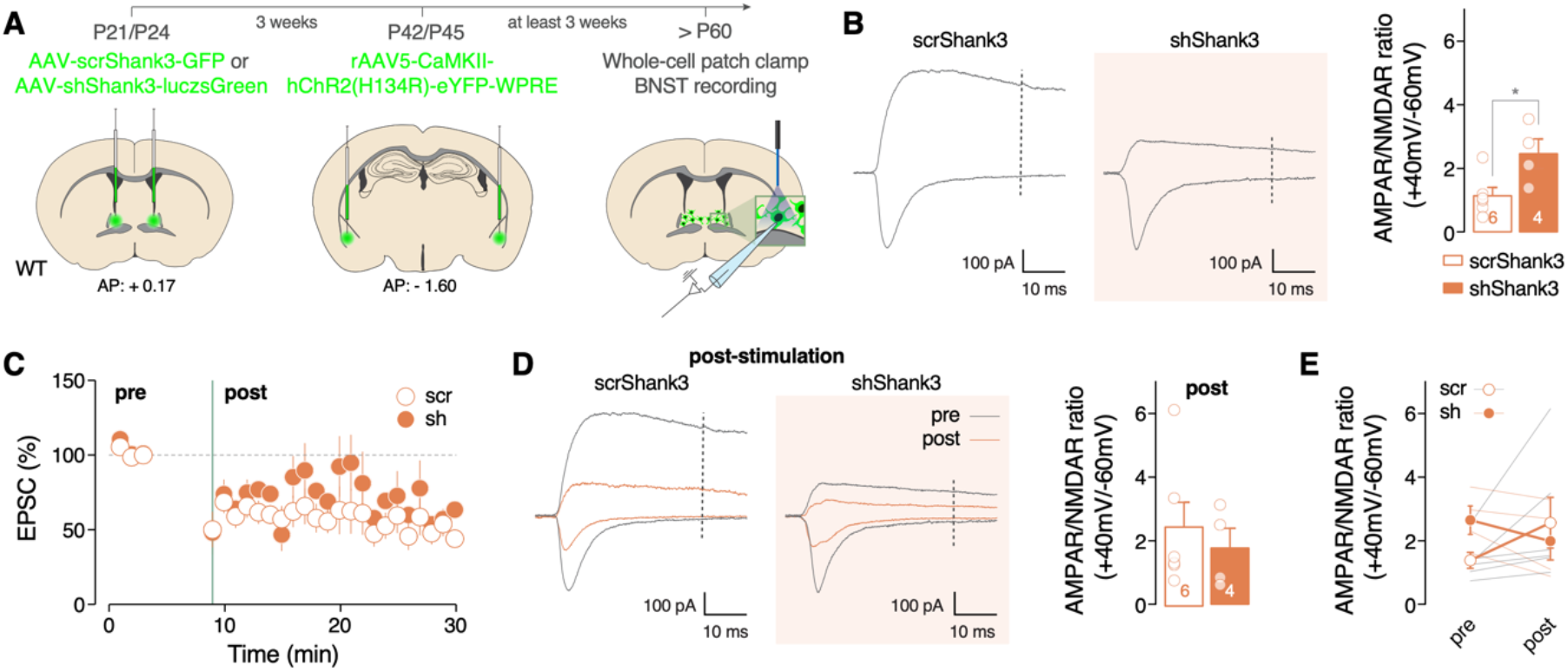
Optogenetic stimulation of BLA to BNST projections elicits long term synaptic plasticity and restore synaptic alterations induced by *Shank3* downregulation. (**A**) Experimental design. (**B**) Example traces and group mean of AMPA/NMDA ratio at BLA to BNST synapses measured at −60 mV and +40 mV for AMPAR- and NMDAR- EPSCs respectively in BNST scr- or shShank3 neurons (unpaired t-test. t_(8)_ = 2.679, P = 0.0280). (**C**) Kinetic of AMPA EPSC amplitude before and after optogenetic stimulation (10 Hz for 5 minutes) for scr and shShank3 positive cells in picrotoxin. (**D**) Example traces (in red) and group mean of AMPA/NMDA ratio at BLA to BNST synapses measured at −60 mV and +40 mV for AMPAR- and NMDAR-EPSCs respectively in BNST scr- or shShank3 neurons after optogenetic stimulation (Mann Whitney test. U = 9, P = 0.6095). (**E**) Comparison of group mean of AMPA/NMDA ratio before and after stimulation for scr- or shShank3 groups (two-way RM ANOVA followed by Bonferroni’s multiple comparisons post-hoc test. Virus x Time interaction: F_(1, 8)_ = 6.246, P = 0.0370; Time main effect: F_(1, 8)_ = 0.5091, P = 0.4958; Virus main effect: F_(1, 8)_ = 0.2044, P = 0.6632). Abbreviations: AP = antero-posterior axis; *p < 0.05.

Until now, we demonstrated that a downregulation of Shank3 in the BNST during adolescence leads to an increased anxious behavior and an altered AMPA/NMDA ratio. As a consequence, we focused our attention in a way to rescue the observed anxiety phenotype. In previous studies, it has been shown that glutamatergic transmission in the BNST plays a crucial role in anxiety control^29,30,38^. Indeed, optogenetic stimulation of the CaMKII terminals coming from the BLA triggered anxiolytic behavior^30^. We therefore examine whether this optogenetic stimulation (10 Hz, 5 ms pulses as reported^30^) could elicit beneficial effects at the synaptic level in our mice with Shank3 deficiency in the BNST. Remarkably, we observed that *ex-vivo* 5 minutes lasting stimulation triggered a LTD in both groups of BNST neurons (**Fig. 3C**). Importantly, we did not find after stimulation a difference in the AMPA/NMDA ratio at BLA to BNST synapses between scr- and shShank3 expressing neurons (**Fig. 3D-E**).

Altogether, these data demonstrated that *Shank3* downregulation in the BNST altered its excitatory synaptic properties, in particular BLA to BNST synapses. Interestingly, induced LTD at that synapses could restore the synaptic alterations provoked by the BNST *Shank3* deficiency.

## Discussion

In the present study, we evaluate the developmental trajectories of social interactions and anxiety like behaviors in *Shank3* deficient mice. We observed that free social interaction alterations emerge at P26 while anxious phenotype appears 10 days after in *Shank3*^*−/−*^ deficient mice. We found that selective *Shank3* downregulation in the BNST during adolescence is sufficient to induce an anxiety phenotype without affecting social interaction and locomotor activity. Associated with these behavioural alterations, we found that Shank3 deficits induced glutamatergic synaptic defects which could be rescued with an optogenetic stimulation of the terminals coming from the BLA.

Exploring *Shank3* deficiency with local gene silencing strategy or with the development of transgenic *Shank3* mouse model gave meaningful insight in the *in vivo* function of Shank3 protein in several brain regions. For example, selective role of *Shank3* has been highlight in the striatum for social interaction and repetitive behaviors, in the VTA for social interactions, in the hippocampus for spatial learning, in the somatosensory cortex in hypersensitivity phenotype^16,24,41,42^. Here, our study pointed to a novel role of *Shank3 in vivo* in the BNST associated with anxiety control. In addition, we stressed the key impact of *Shank3* deficiency in the BNST during adolescence, a crucial time window for anxiety circuit maturation. Here, we focus our study on the impact of BNST *Shank3* deficiency in basal condition. It would be interesting in further studies to evaluate the adaptive responses of BNST *Shank3* deficient mice when exposed to an aversive environment like stressors.

Since Shank3 is a postsynaptic scaffolding protein located at excitatory synapses, we studied the consequences of Shank3 BNST deficiency on glutamatergic transmission. We found at baseline an increase in AMPA/NMDA ratio at BLA to BNST synapses in shShank3 expressing neurons compared to scr expressing neurons. Interestingly, this altered synaptic transmission is rescued *in vivo* by long term depression protocol applied at BLA to BNST synapses. Future studies will determine whether *in vivo* LTD protocol may normalize anxiety deficits induced by selective *Shank3* deficit in the BNST.

Overall, our findings pointed that adolescence is a crucial time window for BNST excitatory synapses maturation and interfering during this developmental stage triggered anxiety trait. These results contributed to a better understanding of the neuronal mechanisms underlying anxiety disorders. Our data proposed a new molecular target essential for the BNST integrity and positioned the BNST as a new player to enrol in diagnosis and future therapeutic developmental strategies during adolescence for anxiety disorders and anxiety disorders implicated in some forms of ASD.

## Authors Contributions

A.C., C.G. and C.B. conceived and designed the experiments. A.C. and C.G. performed and analyzed the behavioural experiments. P.E., and G.C. performed and analyzed the electrophysiological experiments. S.M. performed the RT-qPCR. Y.J. generated the mutated *Shank3* mouse line. A.C., C.G. and C.B. wrote the manuscript and AC prepared the figures.

## Acknowledgments

C.B. is supported by the Swiss National Science Foundation, Pierre Mercier Foundation, ERC consolidator grant and NCCR Synapsy. C.G. is supported by the Foundation for Medical Research: ARF20170938746. P.E. is supported by the Swiss Government Excellence Scholarship (FCS) for PhD studies ESKAS-Nr: 2017.0922. We thank the University of Geneva, the University of Bordeaux and CNRS for the provided infrastructural support. We thank Stamatina Tzanoulinou for her behavioral advice and Lorena Jourdain for technical support.

## Materials and Methods

### Animals

The experimental procedures described here were conducted in accordance with the Swiss laws and previously approved by the Geneva Cantonal Veterinary Authority. Male C57Bl/6j mice were purchased from Charles River and used for scr and sh-Shank3 experiments. C57Bl/6j *Shank3*^Δe4-22^ male and female mice (in this paper *Shank3*^+/+^, *Shank3*^+/−^ and *Shank3*^−/−^) were obtained from Yong-hui Jiang laboratory^16^ and bred in our animal facility. Naive male and female C57Bl/6j were used as stimuli animals in the free social interaction (age and sex were dependent from the experiment). All mice were group-housed (2–5 per cage) in the institutional animal facility under standard 12h/12h light/dark cycles with food and water *ad libitum*. Young male and female mice were weaned and separated by gender at P21. Behavioral experiments were conducted in a room with fixed low illumination (10-15 Lux) and with controlled humidity (40%) and temperature (22-24 °C). The experiments were always performed within a time frame that started approximately 2 hours after the end of the dark circle and ended 2 hours before the start of the next dark circle.

### pViruses and Drugs

Viruses used in this study:

1. Purified scr*Shank3* and sh*Shank3* (AAV1-GFP-U6-scrmbshRNA; titer: 5.9×10^13^ GC/mL and AAV5-ZacF-U6-luczsGreen-sh*Shank3*; titer: 7.4×10^13^ GC/mL, VectorBioLab, sequence targeting exon 21 of the rat and mouse *Shank3* gene: 5′-GGAAGTCACCAGAGGACAAGA-3, as previously described in *Bariselli et al 2016, Bariselli et al 2018*);
2. pAAV-CaMKIIa-eGFP (titer ≥ 3×10^12^ vg/mL, Addgene, catalog # 50469-AAV5) and pAAV5-CaMKII-hChR2(H134R)-eYFP-WPRE (titer ≥ 1×10^13^ vg/mL, Addgene, Catalog # 26969-AAV5).

Drugs used in this study:

1. Picrotoxin (1128, Tocris).

### Stereotaxic viral injections

scr*Shank3* or sh*Shank3* has been injected in the BNST in 3 to 4 weeks old male mice. Anesthesia was induced and maintained with a mixture of oxygen and isoflurane. The animals were then placed on the stereotaxic frame (Stoelting Co, USA) and bilateral craniotomy was made over the BNST at the following stereotaxic coordinates: AP: +0.17 mm, ML: ±0.66 mm; DV: −3.87 mm from bregma. The virus was injected with graduated pipettes (Drummond Scientific Company) at the rate of 100 nL/min for a total of 200 nL per injection site. For all the experiments, the virus was incubated for at least 10 days. AAV5-CAMKII-eYFP (control) and rAAV5-CamkII-hChR2(H134R)-eYFP-WPRE virus were injected bilaterally in the basolateral amygdala (BLA) at the following coordinates: AP:-1.60 mm, ML: ±3.10 mm; DV: −4.00 mm from bregma. The virus was injected with graduated pipettes (Drummond Scientific Company) at the rate of 100 nL/min for a total of 400 nL per injection site. BLA optogenetic experiments were done at least 3 weeks after this injection.

### Open field test

The open field arena was performed in a 40 x 40 x 40 cm arena and a central area of 20 x 20 cm was determined. Each mouse was placed individually in the center of the arena and was filmed for all the duration of the test. For longitudinal study, the male and female mice were allowed to explore the arena for 6 minutes. On the other hand, in the open field test for adult male mice, the task lasted 10 minutes. Distance moved (cm) and time spent in the central area (s) of the arena were measured with the software Ethovision (Noldus, Wageningen, Netherlands). Between each session, the apparatus was cleaned with 70% alcohol.

### Elevated plus maze test

The elevated plus maze consisted of a platform of four opposite arms (40 cm), two of them are open and the other two are closed (enclosed by 15 cm high walls). The apparatus was elevated at 55 cm from the floor. Each male adult mouse was placed individually in the center of the elevated plus maze apparatus with the snout facing one of the open arms and was filmed for 5 min. Distance moved (cm) and time spent in the open and closed arms (s) of the arena were measured with the software Ethovision (Noldus, Wageningen, Netherlands). We calculated the anxiety index as described ^43^:

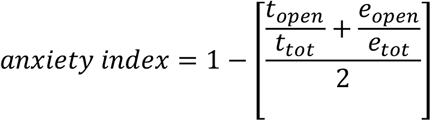

Where *t*_*open*_ =time the animal stayed in the open arms (s)

*t*_*tot*_ =test duration (s) =300 *s*

*e*_*open*_ =number of entries in the open arms (s)

*e*_*tot*_ =total number of entries

Between each session, the apparatus was cleaned with 70% alcohol.

### Free social interaction

The free social interaction was performed in an arena with the same dimension of the homecage of the animals (35 × 20 × 15 cm). Each mouse was placed individually in the center of the arena and filmed during a habituation and a free interaction period. For the longitudinal study, male and female mice were allowed to explore the empty arena for a habitation period of 5 minutes. After this period, a sex-matched adult conspecific was introduced in the arena and free intearction was allowed for 2 minutes. The same social stimulus was used in all the trials of the longitudinal study. In the free social interaction test for adult male mice, sex-matched juvenile (3-4 weeks old) conspecific were used as social stimuli and the period of free interaction lasted 10 minutes. Blind manual scoring was performed in order to quantify the time passed sniffing the social stimulus. Between each session, the apparatus was cleaned with 70% alcohol.

### Histological control

Mice were sacrificed by a transcardiac perfusion with PBS followed by 4% paraformaldehyde diluted in PBS. Brains were post-fixed overnight at 4°C. 50 µm coronal BNST sections were cut and conserved in PBS sodium azide 0.03%. Slices were mounted onto microscope slides with using fluoroshield mounting medium with DAPI (abcam, Cat#ab104139). Images were acquired with an LSM 800 air scan confocal microscope. Mice with wrong injection placement have been removed from this study.

### Real-Time Quantitative Reverse Transcriptase PCR

Similarly to previous studies^41,44^, total RNA was extracted using the RNeasy PLUS Mini Kit (Qiagen), from the BNST of mice infected either with sh*Shank3* or scr*Shank3*. After quantification with Nanodrop Spectrophotometer, 1 μg of RNA was retrotranscribed in cDNA with QuantiTect Reverse Transcription Kit (Qiagen), followed by a PCR amplification of the subsequent transcripts: *Shank3* forward primer 5′-acgaagtgcctgcgtctggac-3′, reverse primer 5′-ctcttgccaaccattctcatcagtg-3′, *Actin* forward primer 5′-agagggaaatcgtgcgtgac-3′, reverse primer 5′ caatagtgatgacctggccgt 3′. Reactions were carried out using iTaqTM Universal SYBR © Green Supermix (Biorad) on Applied Biosystems 7,500 Real-Time PCR System (Applied Biosystems, Foster City, CA, USA) by 50 °C for 2 min, 95 °C for 10 min followed by 40 cycles at 95 °C for 15 s and 60 °C for 1 min. Melting curve analyses were performed to verify the amplification specificity. Relative quantification of gene expression was performed according to the ΔΔ-Ct method^44^.

### *Ex vivo* Electrophysiology

250 µm thick coronal slices containing the bed nucleus of the stria terminalis (BNST) were prepared following the experimental injection protocol described in the text. Slices were kept in artificial cerebrospinal fluid containing 119 mM NaCl, 2.5 mM KCl, 1.3 MgCl2, 2.5 mM CaCl2, 1 mM NaH2PO4 26. 2 mM NaHCO3 and 11 mM glucose, bubbled with 95% O2 and 5 % CO2. Slices were maintained 30 min in bath at 30°C and then at room temperature. Whole-cell voltage clamp recording techniques were used (37°C, 2-3 mL min-1, submerged slices) to measure holding currents and synaptic responses of BNST neurons. The internal solution contained 130 mM CsCl, 4 mM NaCl, 2 mM MgCl2, 1.1 mM EGTA, 5 mM HEPES, 2 mM NA2ATP, 5 mM sodium creatine phosphate, 0.6 mM Na3GTP and 0.1 mM spermine. Currents were amplified, filtered at 5 kHz and digitized at 20 kHz. Access resistance was monitored by a hyperpolarizing step of −4 mV at each sweep, every 10 s. The cells were recorded at access resistance from 10-5 MΩ for BNST neurons. Data were excluded when the resistance changed > 25%. For optogenetic experiments, we stimulated the glutamatergic fibers from the BLA. The stimulus was delivered at 0.1 Hz and the duration was 1–3 ms with a blue LED 485nm. The experiments were conducted in the presence of GABAA receptor antagonist picrotoxin (100 μM); AMPAR-EPSCs were recording at −60mV and measured at the peak, for non-pharmacologically isolated NMDAR EPSCs, we recorded at +40 mV and the NMDA current was measured at 40ms after the stimulation. After this initial AMPA/NMDA ratio we delivered a plasticity protocol that consisted in recordings at −60mV holding potential for 5 min of baseline at 0.1 Hz, followed by 5 min of stimulation at 10 Hz. Then we recorded AMPA EPSCs for 25 min. Then, we measured the AMPA/NMDA ratio as mentioned above. All synaptic responses were collected with a Multiclamp 700 B-amplifier (Axon Instruments, Foster City, CA), and analysed online using Igor pro 6.3 software (Wavemetrics, Lake Oswego, OR).

### Statistical analysis

Statistical analysis was conducted with GraphPad Prism 7 and 8 (San Diego, CA, USA). Statistical outliers were identified with the ROUT method (Q = 1) and excluded from the analysis. The normality of sample distributions was assessed with the Shapiro–Wilk criterion and when violated, non-parametric statistics were applied (Mann-Whitney for two groups comparison, while for multiple comparisons, Kruskal–Wallis or Friedman tests were followed by Dunn’s test). When samples were normally distributed, data were analysed with independent or paired two-tailed sample t-tests, one way, two-way or repeated measures analysis of variance (ANOVA) followed if significant by Bonferroni post hoc tests. All error bars represent the mean ± SEM and the significance was set at p < 0.05. Data were analysed using the graph prism 8 (San Diego, CA, USA).

---

**Sup. Figure 1.**
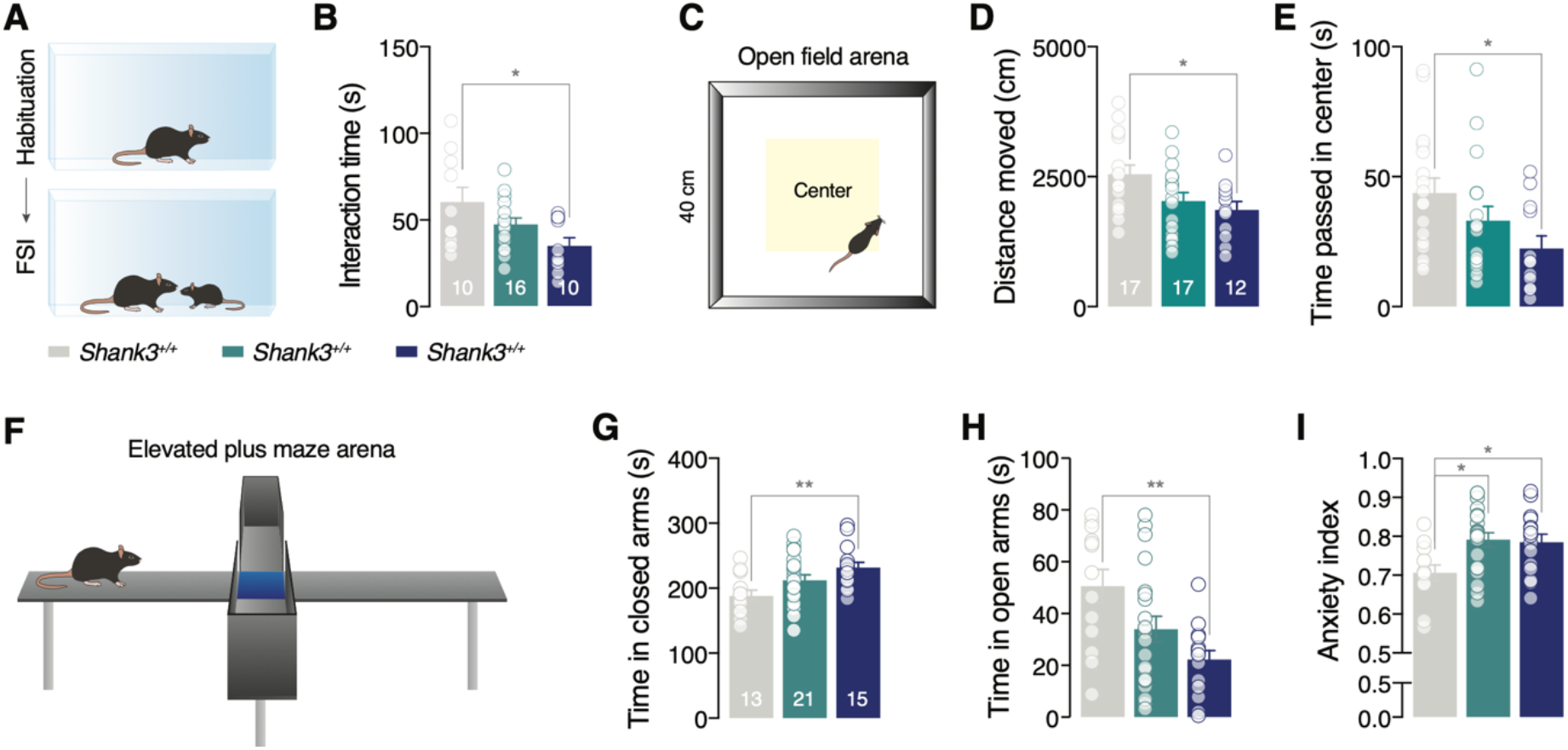
*Shank3*^−/−^ mice present social interaction and exploratory alteration and an anxiety phenotype. (**A, C** and **F**) Experimental schematic. (**B**) Interaction time in free social interaction test for *Shank3*^*+/+*^, *Shank3*^*+/−*^ and *Shank3*^*−/−*^ mice (one-way ANOVA. F_(2, 33)_ = 4.336, P = 0.0213). (**D** and **E**) Distance moved (D: one-way ANOVA. F_(2, 43)_ = 4.340, P = 0.0192) and time passed in the center of the arena (E: Kruskal-Wallis test. P = 0.0280) during the open field for *Shank3*^*+/+*^, *Shank3*^*+/−*^ and *Shank3*^*−/−*^ mice. (**G, H** and **I**) Time passed in the closed (G: one-way ANOVA. F_(2, 46)_ = 5.306, P = 0.0084) and open arms (H: one-way ANOVA. F_(2, 46)_ = 5.148, P = 0.0096) and calculated anxiety index (I: one-way ANOVA. F_(2, 46)_ = 6.595 P = 0.0030) in the elevated plus maze test for *Shank3*^*+/+*^, *Shank3*^*+/−*^ and *Shank3*^*−/−*^ mice. Abbreviations: FSI = free social interaction; *p < 0.05; **0.05 < p < 0.01.

